# Lung megakaryocytes are long-lived, arise from Flt3-negative bone marrow cells, and contribute to platelet recovery in thrombocytopenia

**DOI:** 10.1101/2024.04.11.589077

**Authors:** Alison C. Livada, Kathleen E. McGrath, Michael Malloy, Chen Li, Sara K. Ture, Paul D. Kingsley, Anne D. Koniski, Leah A. Vit, Katherine E. Nolan, Deanne Mickelsen, Grace E. Monette, Preeti Maurya, James Palis, Craig N. Morrell

## Abstract

We previously characterized lung megakaryocytes (Mks) as largely extravascular cells with an immune modulatory phenotype (Pariser et al., 2021). Because bone marrow (BM) Mks are relatively short lived, it is assumed that extravascular lung Mks are constantly ‘seeded’ from the BM, but there are no experimental data to validate this concept. To investigate lung Mk origin and how their origin may impact lung Mk functions, we developed novel models using CFSE dye delivered oropharyngeal (OP) and biotin labeling to specifically label lung Mks and identify lung Mk derived platelets. Labeled lung Mks were present for up to four months, while BM Mks had a less than 1 week lifespan. In a parabiosis model, lung Mks were only partially replaced by a circulating source over a 1-month time period. Unlike tissue resident lung macrophages, we determined using MDS1^-Cre-ERT2^ TdTomato mice that lung Mks arise from a hematopoietic stem cell (HSC) source. However, studies with FlkSwitch mTmG mice showed that lung Mks originate from a Flt3-negative cell lineage, that does not go through a multipotent progenitor stage. CFSE labeling of lung cells enabled us to track lung Mk-derived platelets and we found that about 10% of circulating platelets at steady-state are lung resident Mk-derived, but in the context of sterile thrombocytopenia there was a doubling of lung Mk derived platelets (about 20%). Lung-derived platelets were similarly increased in a murine malaria infection model (*Plasmodium yoelii*) typified by chronic thrombocytopenia. Taken together, our studies indicate that lung Mks arise from a Flt3-negative, HSC-dependent pathway and contribute relatively more platelets during thrombocytopenia.

## Introduction

Platelets serve dual roles as a critical component of thrombus formation and as regulators of immune responses (Morrell et al., 2019). Activated platelets promote inflammation, while circulating resting platelets limit inflammation and maintain vascular integrity (Cloutier et al., 2012; Hilt et al., 2019). Bone marrow (BM) Mks serve as the primary producers of circulating platelets (Asquith et al., 2023; Machlus and Italiano, 2013a), although the presence of Mks in the lung has been described for more than a century (Aschoff, 1893; Kaufman et al., 1965; Levine et al., 1993). Recent endeavors to characterize lung Mks have begun to define their unique phenotype (Livada et al., 2023) and demonstrated that lung Mks contribute to platelet production (Lefrancais et al., 2017), as well as modulating local immune responses (Pariser et al., 2021; Yeung et al., 2020). Better defining the functions and origins of lung Mks will expand our knowledge of Mk roles in vascular biology and provide a better understanding of platelet heterogeneity in health and disease.

While Mks in the lung make platelets, their relative contribution to the total platelet pool remains controversial, with estimates ranging from 7 to 50 % (Howell and Donahue, 1937; Kaufman et al., 1965; Lefrancais et al., 2017). Lefrancais & Ortiz-Muñoz *et al*. estimated the number of platelet fragments generated per lung Mk over time using two-photon intravital microscopy and scaling up these measurements to the full lung volume, they approximated that lung Mks contribute 50% of the platelet pool (Lefrancais et al., 2017). Others have used different imaging methods to extrapolate platelet production potential and proposed that lung Mks contribute fewer numbers of platelets (Asquith et al., 2023). To date, only a few studies have compared lung and BM Mk phenotypes, which have indicated that lung Mks’ roles and functions may be quite different from BM Mks. In addition to platelet production, Mks in the lung have an immune modulatory phenotype (Pariser et al., 2021; Yeung et al., 2020) and can act as antigen presenting cells (Pariser et al., 2021). The lifespan of lung Mks has not been studied, although they are assumed to have a similar life cycle to BM Mks despite the majority of lung Mks being extravascular, 2N, and have functions similar to immune cells. Based on primarily *in vitro* and some *in vivo* data, BM Mks’ life cycle includes maturation, polyploidization, and platelet production, which requires about 5-7 days (d) in humans and 2-3d in rodents, after which they have exhausted their capacity to produce platelets (Liu et al., 2015; Machlus and Italiano, 2013b; Noetzli et al., 2019). Lung Mks are largely 2N and their turnover during homeostasis and pathological states is not known. Additionally, whether lung Mk immune function confers vulnerability, resistance, or heightened responses to challenges has not been studied.

The origin of lung Mks is largely undefined. Lung Mks are assumed to arise from a BM Mk progenitor (MkP) or mature Mks that leaves the BM and travels in the circulation to seed the lung (Lefrançais and Looney, 2019; Lefrancais et al., 2017). Most descriptions of BM megakaryopoiesis describe BM hematopoietic stem cells (HSCs) giving rise to BM Mks in a differentiation process that occurs through successive progenitors with diminishing lineage capacity, until arriving at the megakaryocyte progenitor (MkP) (Boyer et al., 2011; Cheng et al., 2020; Noetzli et al., 2019; Pronk et al., 2007). There is also a less defined ‘direct’ pathway from HSC to MkP or Mk (Haas et al., 2015; Rodriguez-Fraticelli et al., 2018; Sanjuan-Pla et al., 2013; Shin et al., 2014; Yamamoto et al., 2013) that may be particularly salient during immune challenge (Haas et al., 2015). The lineage of lung Mks and whether they arise from successive progenitors or directly from an HSC either locally or from the BM is not known.

We now demonstrate that lung Mks are long-lived cells that contribute about 10% of the circulating platelet pool at steady state and increase their relative platelet production in the setting of thrombocytopenia. Using lineage tracing reporter mice, we found that lung Mks are an HSC-dependent population, but arise via a Flt3-independent lineage from HSCs, in contrast to largely Flt3-dependent differentiation for BM Mks. These data indicate a distinct lineage for lung Mks and suggest that the Mk origin may confer lung Mks with a longer lifespan and greater contribution to low platelet states. This functionally indicates that resident lung Mks are a stable source of platelets in times of increased demand.

## Results

### Lung Mks are long-lived cells

To begin to investigate the function and lifespan of lung Mks, we modified protocols used to track immune cells (Jenkins et al., 2021; Legge and Braciale, 2003) and developed a novel strategy to specifically label lung Mks. We administered CFSE dye via an oropharyngeal (OP) route to label lung cells, but not cells in other tissue beds (Fig 1A). CFSE is a cell-permeable, fixable dye that allows for long-term and stable cell labeling. Six hours after administering CFSE dye OP to wildtype (WT) mice, we found that lung Mks were CFSE^+^ (Figure 1A: bottom row), whereas BM (Figure 1A: top row) and spleen (Figure 1A: middle row) Mks were not. To determine a time course for lung Mk labeling, we repeated the CFSE dye delivery and determined the presence of CFSE^+^ Mks on day one (d1), d3, d5, and d7 post-CFSE OP administration. There were negligible CFSE^+^ Mks in the BM and spleen, while a substantial number of CFSE^+^ Mks remained in the lung over the 7d timeframe (Figure 1B). This method also labeled Lineage^-^, c-Kit^+^ (LK) cells in the lung, but not BM or spleen (Supplemental Figure 1A). Therefore, delivery of CFSE OP yielded tissue-specific labeling of lung Mks and LKs, with no spillover to the BM or spleen. As a complementary approach, we labeled cells with EZ-Link-NHS-Biotin, (referred hereafter as biotin label) (Nygren and Bryder, 2008). Biotin was administered OP on d0 to WT mice and streptavidin binding used to identify biotin labeled cells. On d5 post-OP biotin delivery, we found that about 40% of lung Mks were streptavidin^+^, whereas no streptavidin binding was detected in BM or spleen Mks (Figure 1C). These data indicated that at least a population of lung Mks live in the lung for an extended time. To assess the long-term lifespan of lung Mks, we gave CFSE OP to mice and waited until d30 or d120 to harvest BM, lungs, and spleens. Surprisingly, a significant number of lung Mks remained CFSE^+^ on both d30 and d120 post-labeling, while no CFSE^+^ BM and spleen Mks were detected (Figure 1D). Hematopoietic progenitors, LK cells, also remained CFSE^+^ only in lung tissue at d30 and d120 (Supplemental Figure 1B). These data imply that at least some lung Mks remained quiescent and long-lived cells or arose from a local CFSE^+^ progenitor.

**Figure 1.**
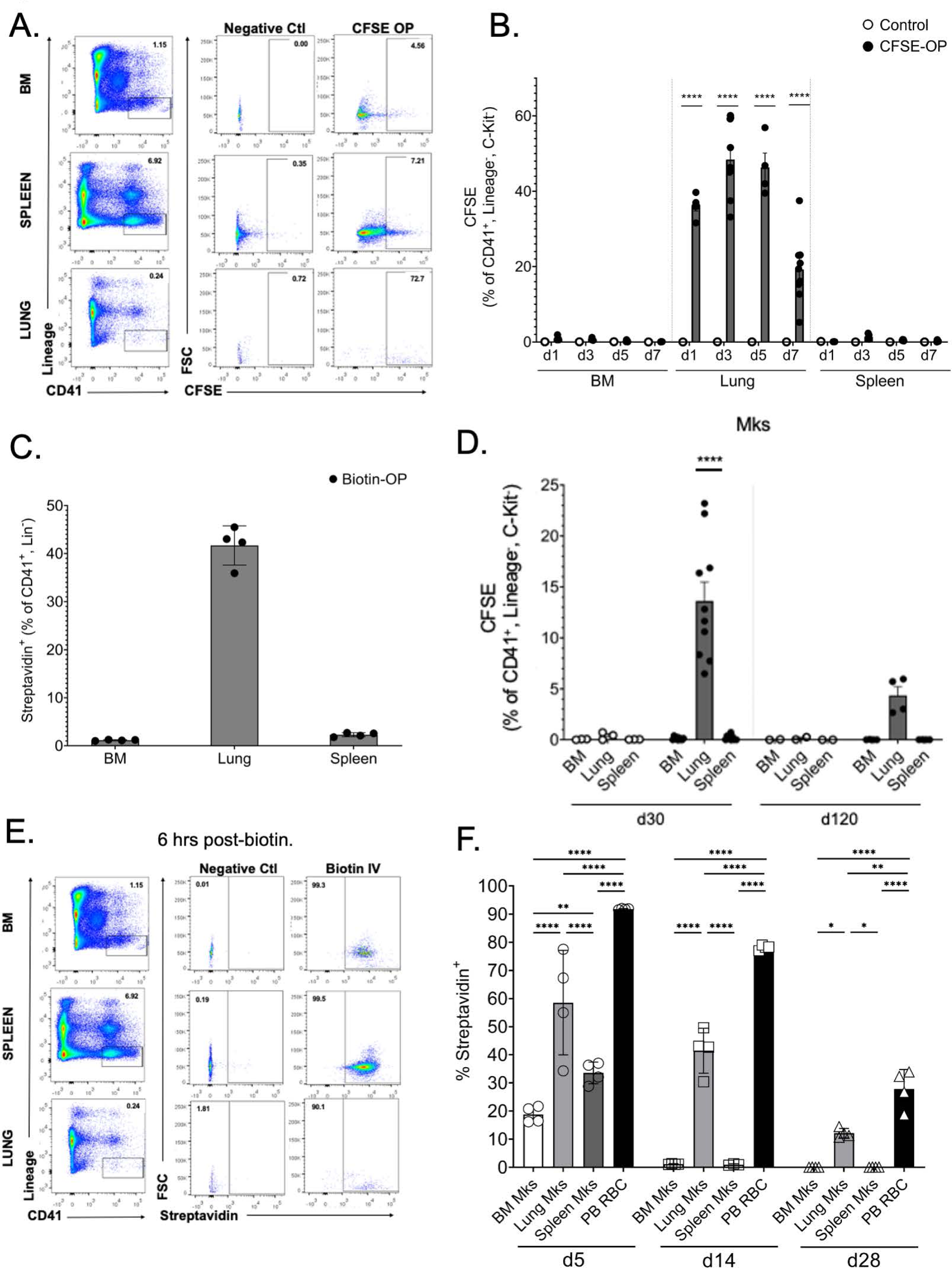
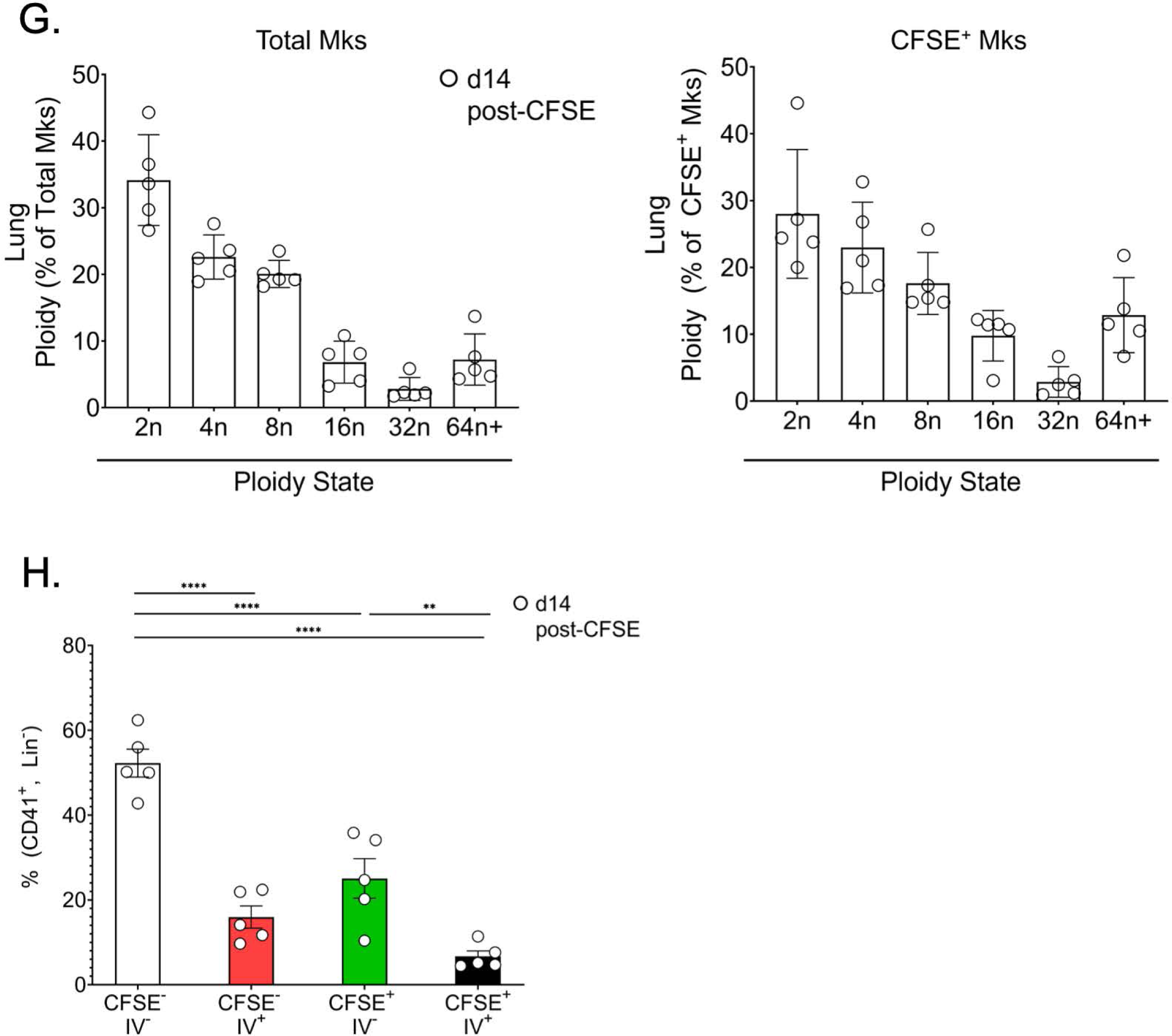
Lung Megakaryocytes are Long-Lived Cells. A) CFSE delivered OP to mice specifically labeled lung, but not BM or spleen Mks. OP delivery of B) CFSE or C) biotin labeled lung Mks, but not Mks in the BM or spleen up to 7d after *in vivo* labeling. D) CFSE labeled lung Mks are present for up to 120d after *in vivo* OP delivery. E) Biotin similarly labeled Mks in the BM, spleen, and lung 6 hrs after given to mice IV. F) Biotin labeled lung, but not BM and spleen Mks were present up to 28d following IV delivery to mice. RBCs as positive control. G) Total lung Mks and lung resident CFSE^+^ Mks have similar ploidy (d14 post-CFSE). H) CFSE^-^ and CFSE^+^ lung Mks have a similar intra- and extravascular distribution. (\**p*-value < 0.05, ± SEM; B), D), and F) Two-Way ANOVA with Tukey’s multiple comparison correction tests; H) One-Way ANOVA with Sidak multiple comparison correction tests).

To provide a comparative analysis of Mk tissue residence, we gave biotin to mice intravenously (IV) to label all tissue compartments and waited 6 hrs to determine the extent of Mk labeling in the BM, spleen, and lung. Representative gating demonstrated >90% labeling of Mks in each tissue (Figure 1E). We next determined streptavidin binding^+^ Mks on d5, d14, and d28 post-biotin IV delivery. Circulating red blood cells (RBCs) were used as a positive control (90% labeled on d5), and as expected, remained 75% and 30% biotin^+^ on d14 and d28, respectively. About 20% of BM Mks remained streptavidin^+^ on d5, but none were noted after d5 (Figure 1F). Spleen Mks remained 30% streptavidin^+^ on d5, but like BM Mks, none were noted at the later time points. In contrast, streptavidin^+^ lung Mks were present at all time points tested (41% and 12% on d14 and d28, respectively) (Figure 1F). These data demonstrated that at least a substantial number of lung Mks have a much longer tissue residence compared to BM Mks.

To further characterize the CFSE^+^ lung Mks, we evaluated their ploidy state, and intravascular vs extravascular location on d14 post-CFSE administration. Similar to our prior report (Pariser et al., 2021), total lung Mks were largely low ploidy (<4N, Figure 1G, left panel). CFSE^+^ Mks were also largely low ploidy with a similar range of ploidy states (Figure 1G, right panel). At this time point, we also found that about 25% of total Mks were CFSE^+^ extravascular and about 7% were CFSE^+^ intravascular (Figure 1H), similar to the overall 3:1 extravascular:intravascular ratio previously reported (Pariser et al., 2021). CFSE^+^ Mks are therefore primarily low ploidy and exist in both the extravascular and intravascular space.

Taken together, these labeling studies indicate that at least a subset of lung Mks are long-lived and maintain themselves locally for up to 120d.

### Lung Mks are only partially replaced by a BM source

To determine whether long-lived Mks are tissue-resident, we used a parabiosis mouse model. WT and mTmG mice were given CFSE OP to label lung cells a week prior to parabiosis surgery. Mice were joined for one-month, after which blood, BM, lungs and spleens were assessed for parabiont chimerism and CFSE^+^ lung cells (Figure 2A). Alveolar macrophages (AMs) are tissue-resident and served as a positive control (Hashimoto et al., 2013), whereas interstitial macrophages (IMs) are circulatory derived and served as a negative control for tissue residence (Chakarov et al., 2019). One month after parabiosis surgery, all AMs remained CFSE^+^, host-derived (green, Figure 2B: left top panel). As expected, the vast majority (77%) of IMs were replaced by CFSE^-^ host-derived cells (white, Figure 2B: right top panel). After one month of parabiosis, 23% of the IMs were CFSE^-^ partner derived (red, Figure 2B: right top panel), indicating circulatory replacement of IMs from the partner parabiont. No IMs were CFSE^+^ from the partner (orange) or host (green). These results demonstrated that parabiosis yielded circulatory mixing and confirmed the CFSE labeling of tissue-resident cells. When we evaluated lung Mks, we found an intermediate phenotype between IMs and AMs (Figure 2B: bottom panel). Like IMs, more than half (63%) of lung MKs were replaced by CFSE^-^ host-derived cells (white) in the 1-month time-period. About 19% of lung Mks were CFSE^-^ partner-derived (red), and none were CFSE^+^ partner-derived (orange). However, unlike IMs, there was a subset (∼15-20%) of lung Mks that remained CFSE^+^ host-derived (green). These experiments indicated that over a month time period, the circulation contributed a portion of lung Mks, but also that a population of lung Mks had a longer lifespan and did not undergo circulatory replacement.

**Figure 2.**
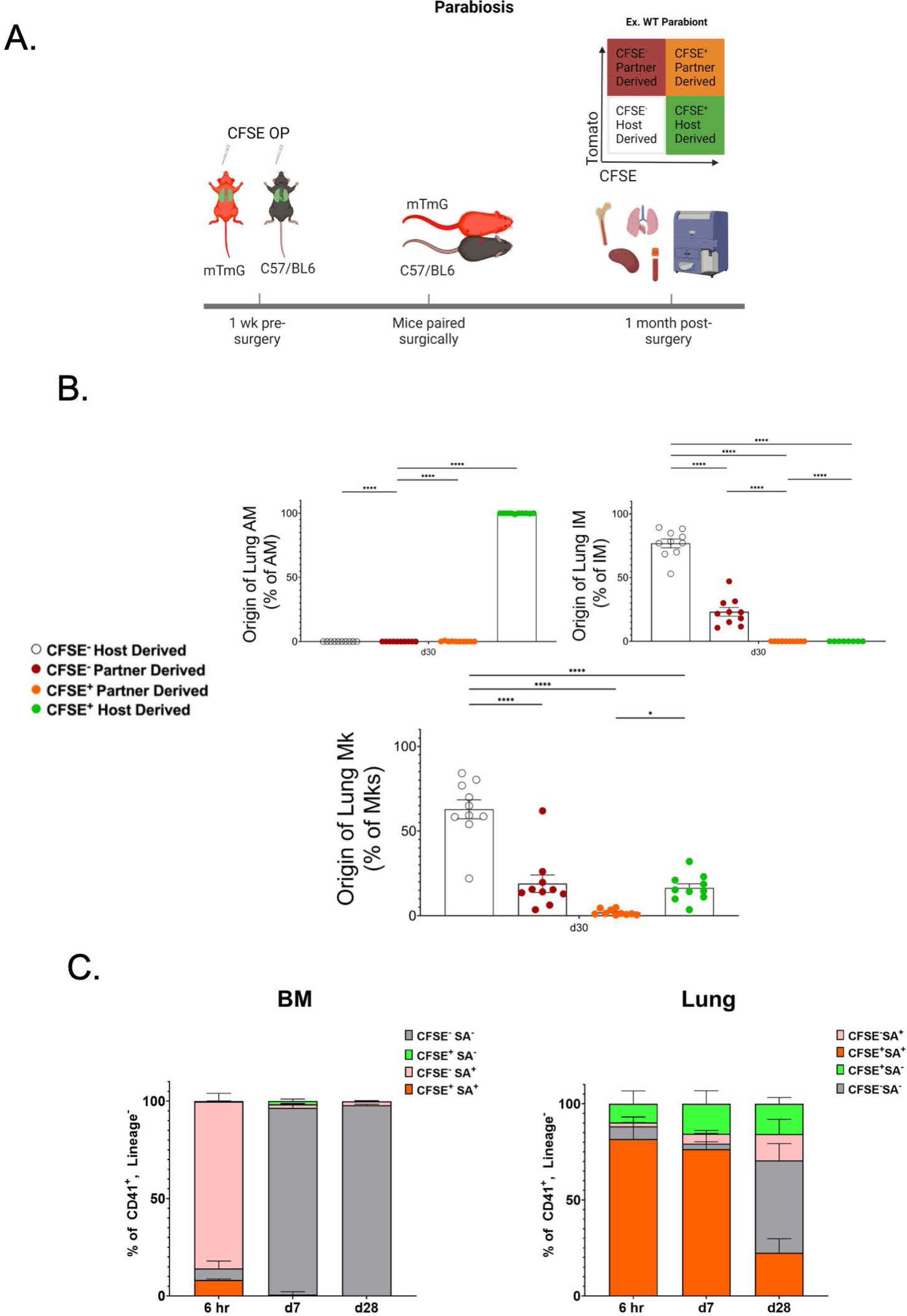
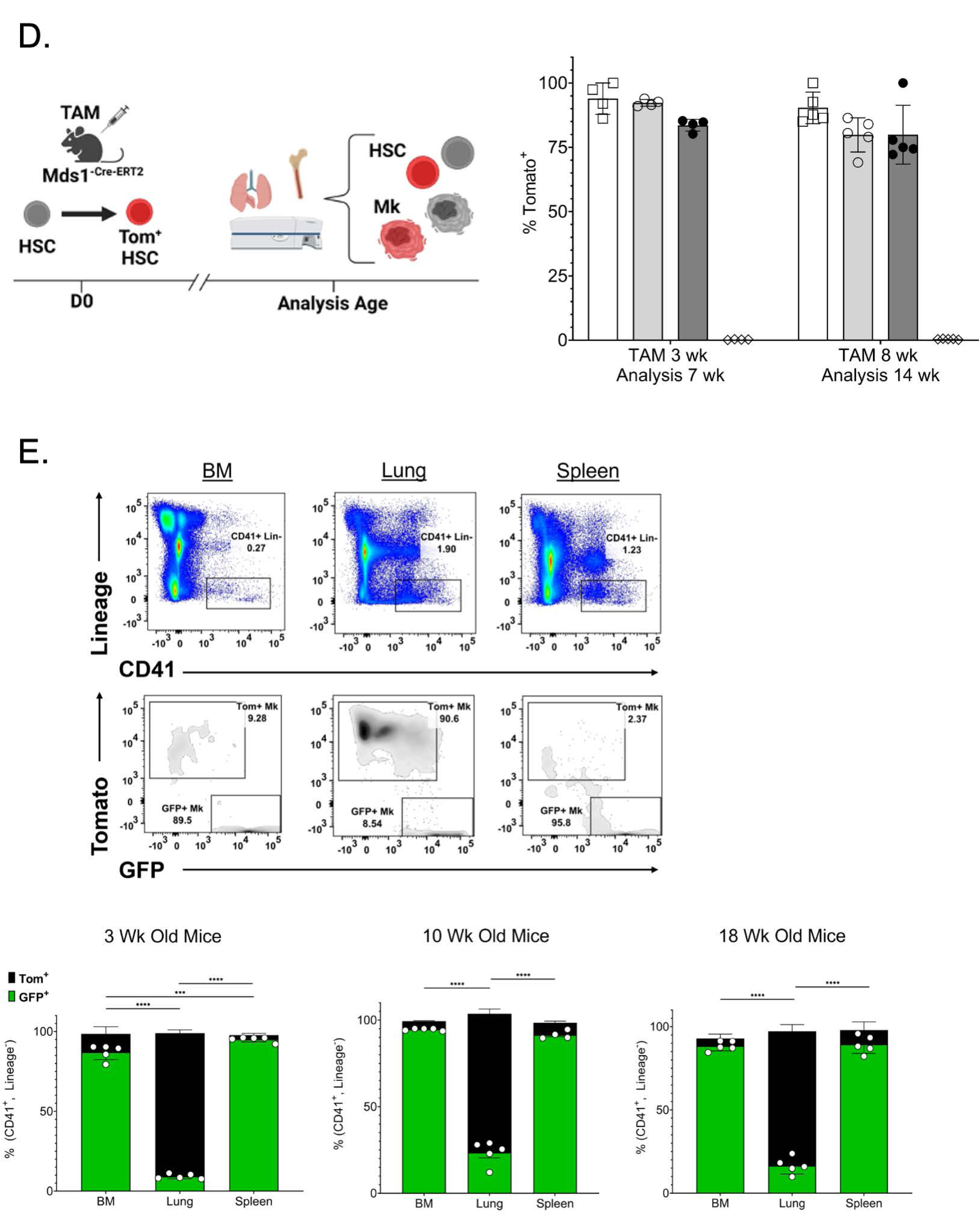
Lung Megakaryocytes are Bone Marrow Derived. A) mTmG and C57/BL6 mice were given CFSE OP and 1-wk later parabiosis surgeries performed. B) 1 month later, lung alveolar/resident macrophages (AM), interstitial macrophages (IM) and lung Mk sources were determined. As expected, AMs were tissue resident and IMs were circulatory source derived. Lung Mks were only partially replaced by a BM source. C) Mice were given CFSE OP and biotin IV. 6 hrs, 7d and 28d later BM and lung Mks were assessed. BM Mks were replaced within 7d, but lung resident Mks were found at each time point tested. D) MDS1-TdTomato^Cre-ERT2^ mice were treated with Tamoxifen to label HSCs and HSC derived cells. Lung Mks were HSC derived. E) BM, spleen and lung Mks in FlkSwitch mice were assessed at multiple ages to determine whether Mks were derived in a Flt3-dependent differentiation pathway. Lung Mks largely remained Tom^+^/Flt3^-^ indicating a potential direct HSC to Mk differentiation that is distinct from BM and spleen Flt3 dependent differentiation. (\**p*-value < 0.05, ± SEM; B) One-Way ANOVA with Sidak multiple comparison correction tests; D) and E) Two-Way ANOVA with Tukey’s multiple comparison correction tests)

To provide a complementary approach and confirm the data indicating the presence of a subset of lung Mks that has a longer tissue-resident lifespan, we combined the CFSE OP and biotin IV labeling methods. Mice received CFSE OP and biotin IV on d0, and BM and lung Mks were assessed 6 hrs later, on d7 and on d28 for CFSE dye and biotin labeling. At the 6 hr time point, BM Mks were largely (>90%) CFSE^-^, streptavidin^+^ shown in pink bar (Figure 2C: left panel). On d7 and d28, BM Mks did not retain biotin labeling, and were almost completely double negative (grey bar). The lung was largely double positive at 6 hrs (orange bar) and remained so on d7 (Figure 2C: right panel). On d28, about half the lung was double negative for CFSE and streptavidin, while the other half retained either dye, biotin, or both. These results are very similar to the parabiosis model and indicate that a significant number of lung Mks are tissue resident for over 30d.

### HSCs give rise to lung MKs via a Flt-3-independent pathway

Given this intriguing discovery of long-lived lung Mks that are replaced by a circulating source, we sought to better define their origin. Lung resident macrophages are initially seeded from an HSC-independent source during embryo development. We therefore leveraged a lineage tracing model to track HSC-dependent hematopoiesis, a MDS1^-Cre-ERT2^ TdTomato mouse model (Zhang et al., 2021). After induction with tamoxifen TdTomato is expressed in HSCs and the fluorescent reporter is retained in the subsequent HSC daughter cells (Figure 2D). Tamoxifen administration at 3 wks of age revealed close to 100% Tomato labeling of BM LT-HSCs collected at 7 wks (Figure 2D). As a negative control, we used brain microglia, a tissue-resident, HSC-independent cell derived from the yolk sac during embryogenesis (McGrath et al., 2015), and found 0.23% Tomato labeling of microglia. 90% of BM Mks and 85% of lung Mks were Tomato-labeled (Figure 2D). Similarly, when we administered tamoxifen at 8 wks and collected tissues at 14 wks, lung and BM Mks were Tomato-labeled while microglia were not (Figure 2D). These data indicated that from early post-natal and through adult life, lung Mks are HSC derived.

Some Mks arise directly from HSCs, but not through a MPP stage, a process sometimes termed emergency megakaryopoiesis (Carrelha et al., 2018; Haas et al., 2015; Sanjuan-Pla et al., 2013; Yamamoto et al., 2013). To more specifically define the source of HSC-derived lung Mks, we used the FlkSwitch-mTmG model (Boyer et al., 2011) in which cells are Tomato^+^ unless a cell turns on Flt3-protein expression, excises Tomato, and then express GFP. Flt3 is expressed during the transition to short-term HSCs (ST-HSCs) and is highly expressed in multipotent progenitors (MPPs) and lymphoid cells (Beaudin et al., 2016; Boyer et al., 2011). Because the recombinase alters the genome, Flt3-expressing progenitors yield daughter cells that remain GFP^+^. As expected in 3 wk, 10 wk and 18 wk old FlkSwitch-mTmG mice, BM and spleen Mks were largely GFP^+^ (representative gating and quantification, Figure 2E). In contrast, lung Mks were Tomato^+^ (Figure 2E). We also confirmed that the lung Mk Tomato expression was not due to cleavage of GFP or other tissue-preparation related issues, as lung interstitial macrophages were largely GFP^+^ and alveolar macrophages were Tomato^+^ (Supplemental Figure 2A). Together the FlkSwitch and MDS1 models suggest that lung Mks are derived from HSCs that differentiate via a Flt3-independent progenitor pathway.

### Lung-resident Mk derived platelets make up a minor portion of the circulating platelet pool at steady-state

The literature has described a pathway from Flt3^-^ LT-HSCs directly to a Mk or Mk progenitor (MkP), particularly in the setting of emergency thrombopoiesis (Carrelha et al., 2018; Haas et al., 2015; Nishikii et al., 2015; Sanjuan-Pla et al., 2013; Yamamoto et al., 2013). We therefore sought to assess the platelet production response of lung Mks during both basal and thrombocytopenic states. To quantify resident lung Mk-derived platelets at steady state, we isolated platelets from mice given CFSE OP and CD42C-X649 antibody IV to label all lung-derived platelets (CFSE^+^) and platelets present on d0 (X649^+^). Using flow cytometry, we selected CD41^+^ events and evaluated CFSE against d0 CD42C-labeling antibody (Figure 3A). On d1 post-CFSE, the vast majority of CD42C^+^ platelets labeled on d0 were also CFSE^+^ (Figure 3A: third column), attributable to some leak of CFSE dye into the vasculature that does not reach tissue compartments. By d5 post-CFSE, we found a small number of new CD42C^-^/CFSE^+^ platelets, and no longer saw significant numbers of CD42C^+^/CFSE^+^ platelets (Figure 3A: fourth column). The number of new CFSE^+^ platelets as a percent of total platelets ranged from 1-6% on d3 and d5 post-CFSE (Figure 3B: left panel). Because we did not have full labeling of all lung Mks (Figure 1B), we normalized CFSE^+^ platelets to CFSE^+^ Mks to represent the maximum potential contribution of lung resident Mks to platelet production. This normalization suggests that about a maximum of 10% of circulating platelets are lung-resident Mk derived (Figure 3B: right panel). We used a complementary biotin labeling OP approach to identify streptavidin binding^+^ platelets derived from lung Mks. Similar to the CFSE model, about 5% of platelets were streptavidin^+^ on d5 post-biotin OP administration (Figure 3C). Taken together, these data indicated that at steady state about 5-10% of circulating platelets come from lung resident Mks.

**Figure 3.**
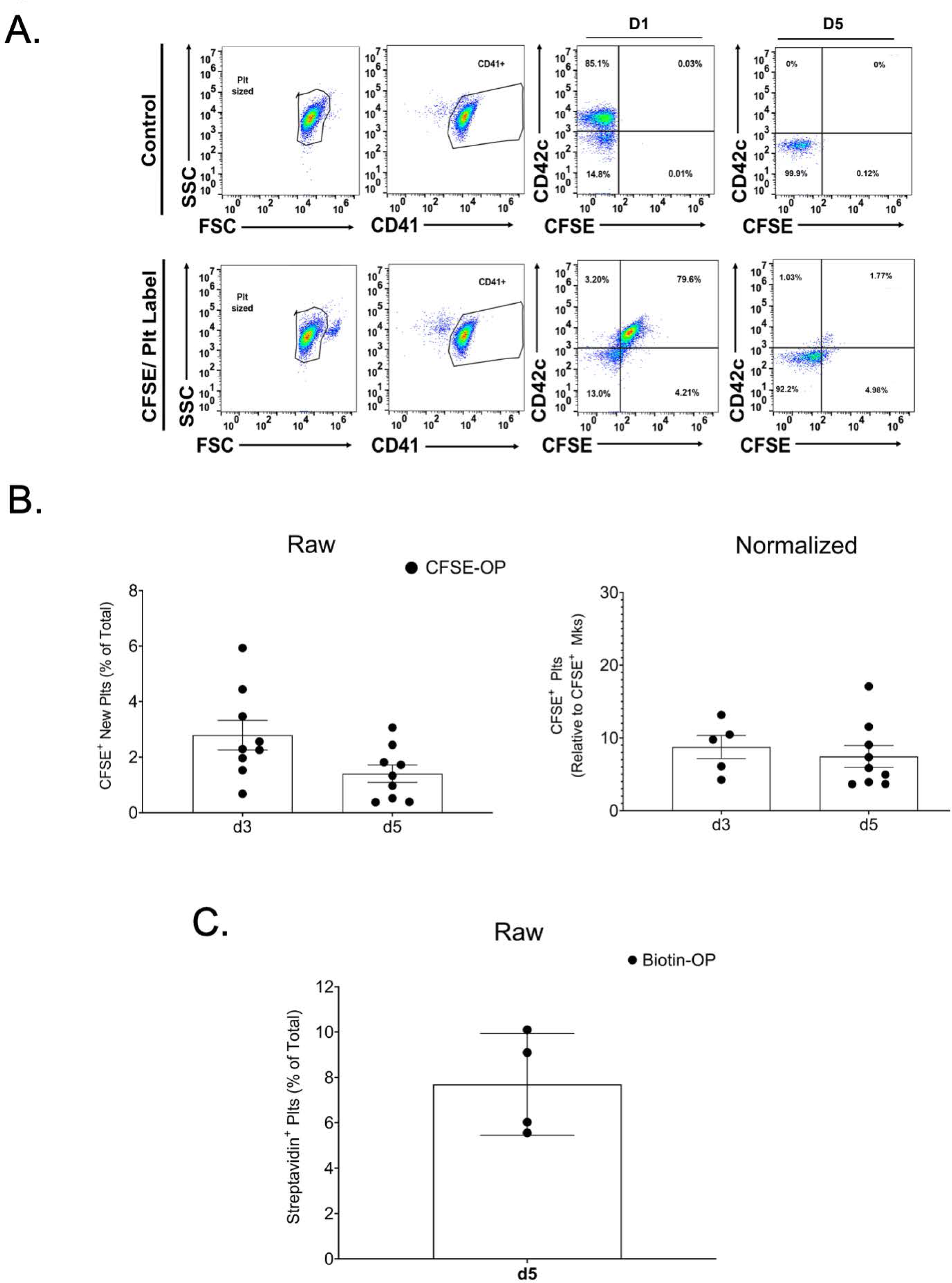
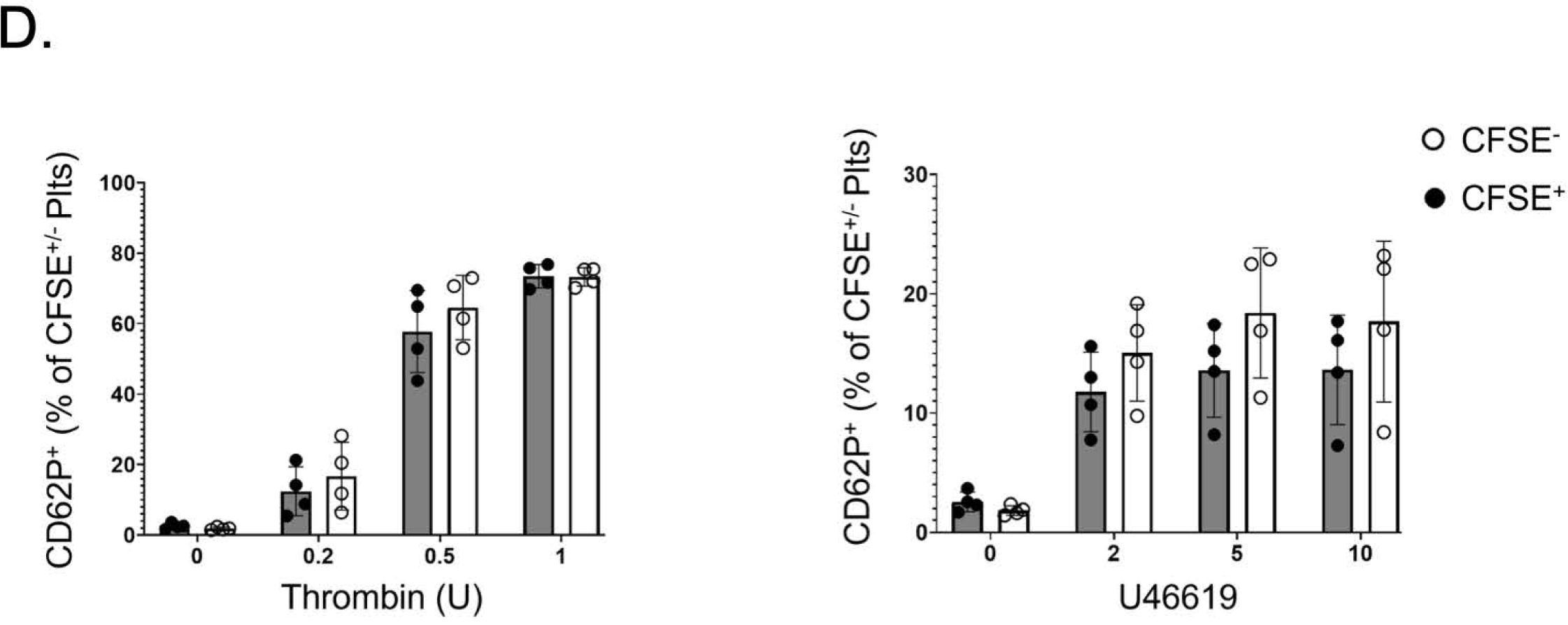
Lung Resident Megakaryocytes Produce Platelets. A) Mice were simultaneously given CFSE OP and anti-CD42c platelet labeling antibody IV. Lung derived platelets were quantified as CFSE^+^/CD42c^-^ platelets by flow cytometry. B) 2-5% of total platelets were lung derived, and when normalized to CFSE^+^ Mks, lung derived platelets represented a maximum of about 10% being lung resident Mk derived. C) CFSE data were validated by biotin delivered OP and quantifying strepavadin binding platelets. D) CFSE^+^ lung derived and CFSE^-^ platelets were agonist stimulated and platelet activation quantified by flow cytometry for CD62P surface expression. Lung derived platelets and marrow derived platelets responded similarly in response to thrombin and U46619. (\**p*-value < 0.05, ± SEM; D) Multiple *t*-tests with Holm-Sidak multiple comparison correction test)

Next, we assessed lung-derived platelet reactivity to common agonists. Platelets were stimulated with either thrombin (Figure 3D: left panel) or the thromboxane receptor agonist U46619 (Figure 3D: right panel) and platelet stimulation measured using CD62P^+^ surface expression. Similar agonist responses occurred at every dose tested for both CFSE^-^ and CFSE^+^ platelets (Figure 3D). From these experiments, we concluded that lung resident Mks produce about 10% of the platelet pool during homeostatic conditions, and these platelets reacted similarly to common agonists like thrombin and thromboxane mimetic.

### Lung-derived platelet production increases with acute and chronic thrombocytopenia

Our prior studies, and those of others, demonstrated that lung Mks are immune differentiated (Lefrancais & Ortiz Munoz et al, 2017; Pariser et al, 2021; Yeung et al 2021). We therefore evaluated the role of lung Mk platelet production in inflammatory contexts by testing the effect of LPS. Mice were simultaneously given CFSE OP and LPS (1 mg/kg) or PBS intraperitoneal (IP). 24 hrs later, platelet counts were significantly reduced in the LPS treated mice (Figure 4A: left top panel). 3d later, CFSE^+^ lung-derived platelets as a percent of total platelets and normalized to CFSE^+^ Mks were increased (p value = 0.0597; Figure 4A, right panels). No significant changes in Mk numbers were observed in BM or lungs on d1 or d3 post-LPS (Figure 4A, left bottom panel). To determine whether an inflammatory signal without associated thrombocytopenia increased lung Mk production, we treated mice with IFNγ. We saw no change in total or CFSE^+^ lung-derived platelets (Figure 4B), indicating that platelet production in the lung is not changed in the setting of immune stimuli without concomitant thrombocytopenia.

**Figure 4.**
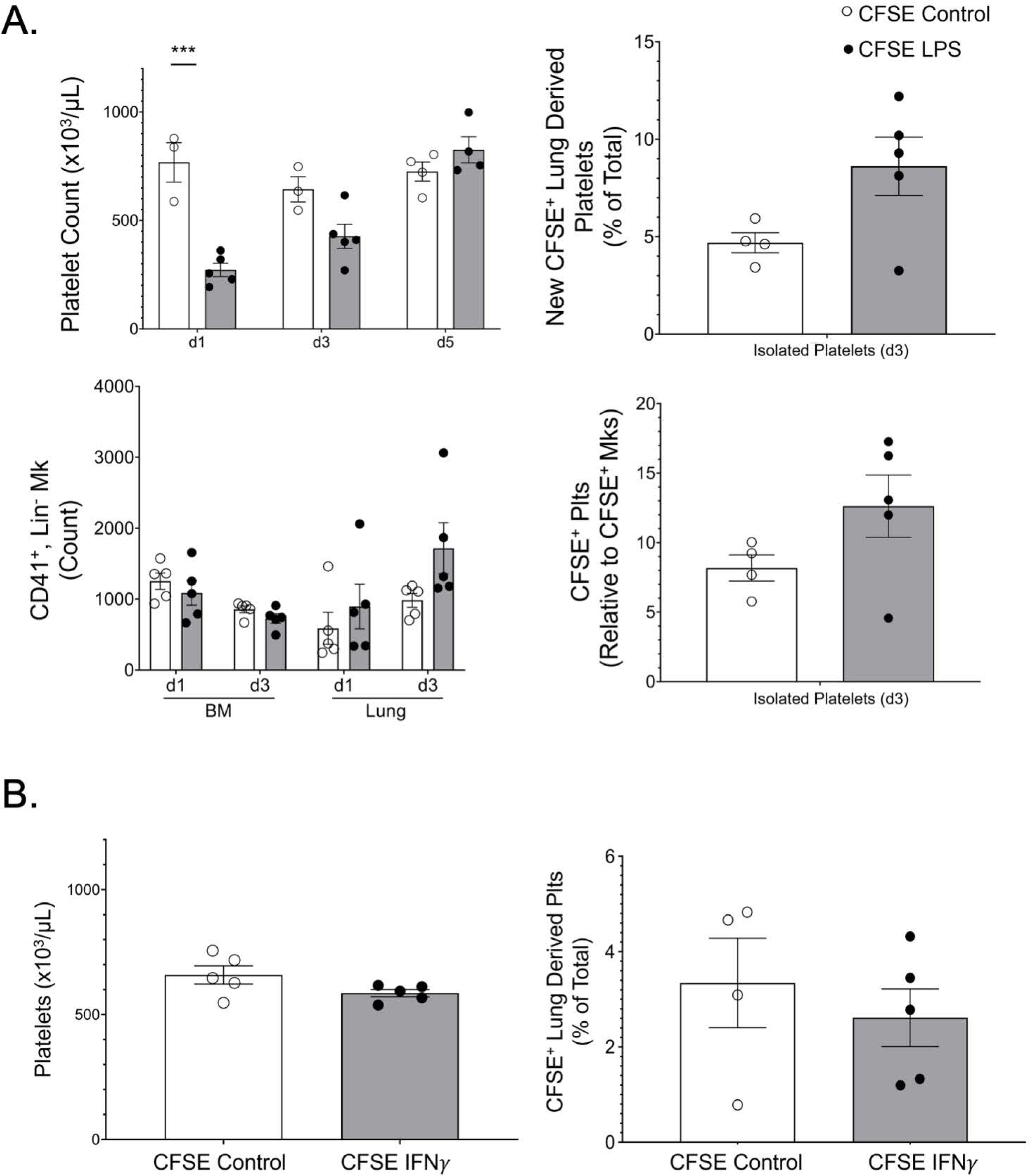
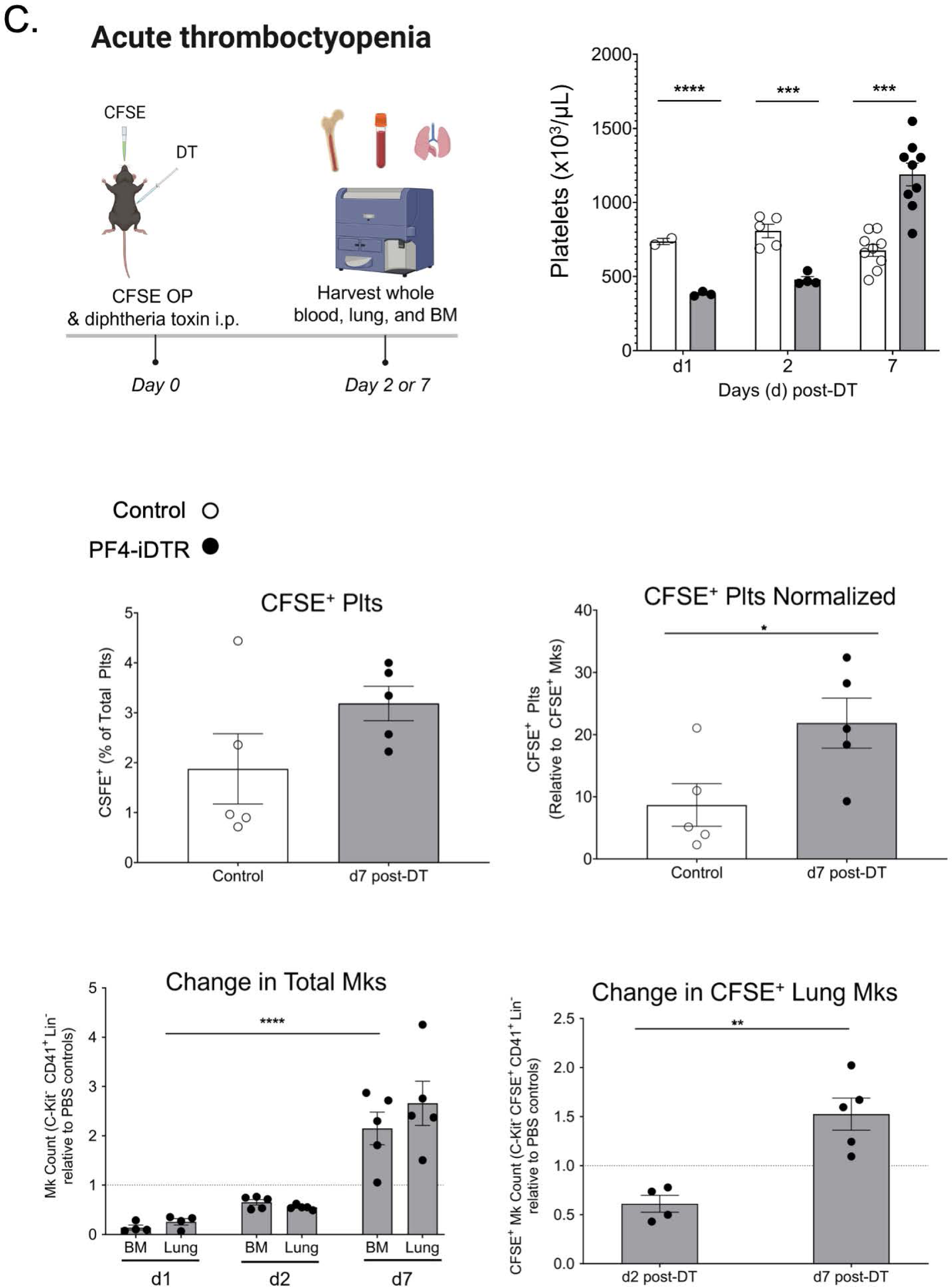
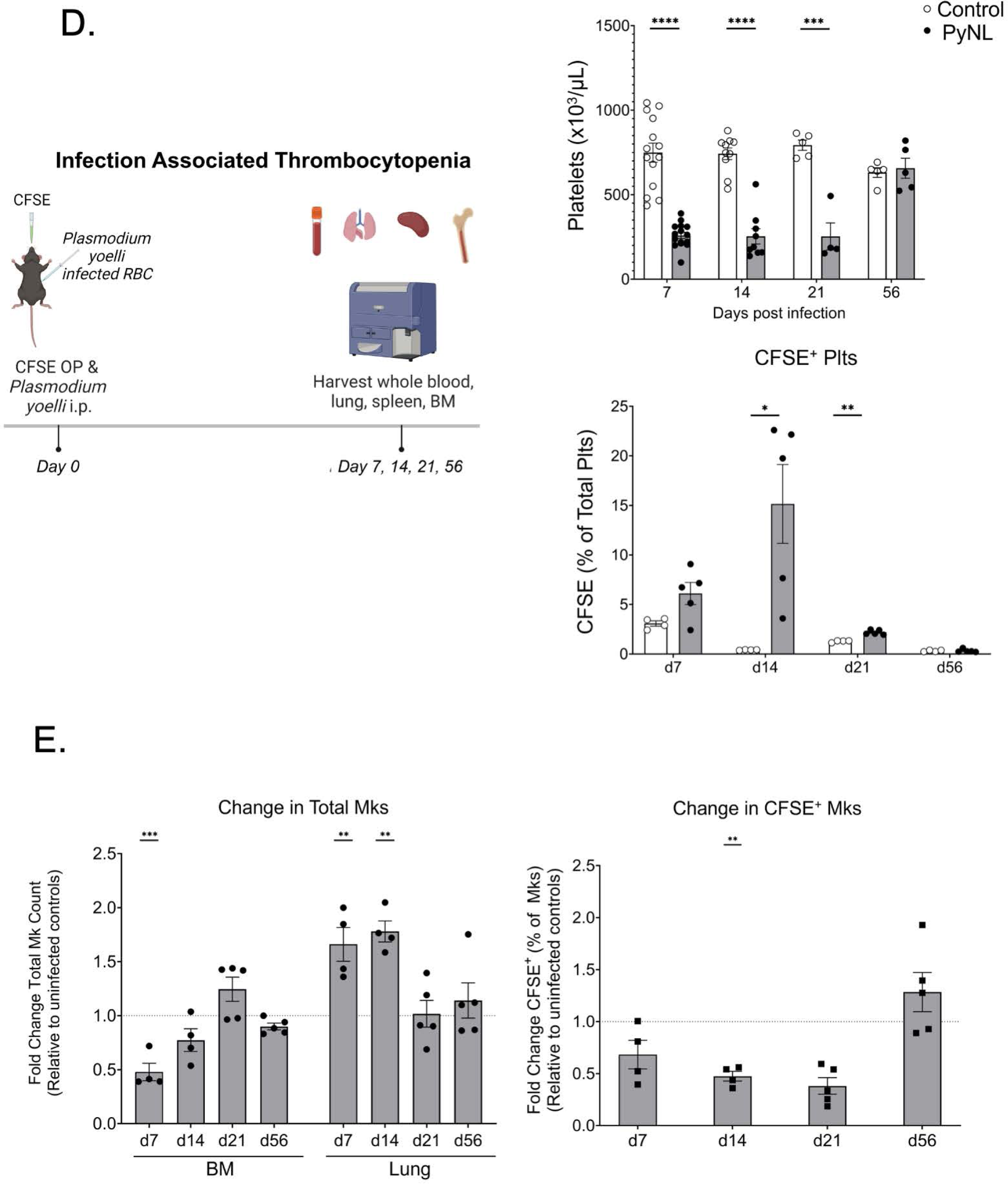
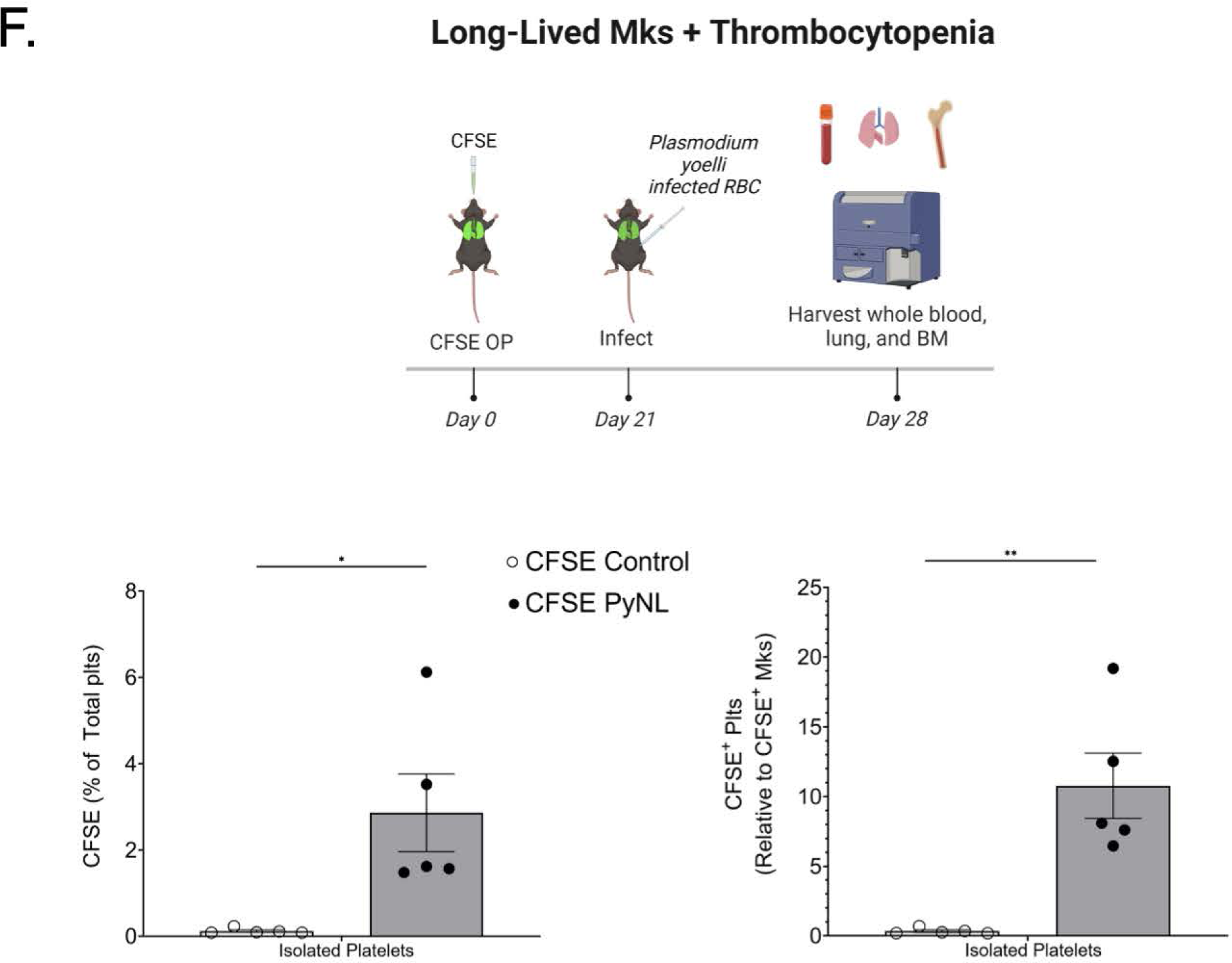
Lung Resident Megakaryocytes Respond to Thrombocytopenia. A) Mice were given CFSE OP and LPS or PBS IP. LPS reduced platelet, but not Mk counts, and increased lung derived platelets. B) IFNγ had no effect on platelet counts or CFSE^+^ platelets. C) PF4^cre^-iDTR mice treated with DT had reduced platelet counts and increased CFSE^+^ lung derived platelets. DT treatment reduced BM and lung total Mks as well as CFSE^+^ Mks on d2 post-DT. CFSE^+^ Mks relative to total Mks in control mice increased on d7 post-DT when Mk and platelet counts were recovered. D) Lung resident Mk derived platelets were increased with chronic infection associated thrombocytopenia. Mice infected with PYnL had reduced platelet counts and more lung derived, CFSE^+^ platelets in the first 2 weeks post-infection. E) Following PYnL infection, the number of BM Mks declined over the first 2 wks, but the number of lung Mks was increased. The proportion that were CFSE^+^ was reduced early post infection, indicating an influx of new Mks from an extra-pulmonary source. After infection recovery the relative per cent of CFSE^+^ Mks in the lung were similar to control mice at the same time post-CFSE, indicating an expansion of lung resident Mks from a source present at the time of infection. F) Lung resident Mks can make platelets. Mice were treated with CFSE OP and 3 wks later infected with PYnL. CFSE^+^ platelets were increased 1 wk post-infection (4 wks post-CFSE). (\**p*-value < 0.05, ± SEM; A) and C) Platelet count: Multiple *t*-tests with Holm-Sidak multiple comparison correction test; CFSE % and CFSE normalized: unpaired two-tailed *t*-test; Mks: Two-Way ANOVA with Tukey multiple comparison correction tests; B) and F) unpaired two-tailed *t*-tests; D) Multiple *t*-tests with Holm-Sidak multiple comparison correction test; E) Two-Way ANOVA with Sidak multiple comparison correction tests).

Based on these data, we sought to determine whether inflammation was required to induce increased lung-derived platelet production in an acute thrombocytopenia context. PF4-iDTR mice (Li et al., 2024; Wuescher et al., 2015) were administered diphtheria toxin (DT) to induce Mk apoptosis and acute thrombocytopenia for up to 5d. On d0, CFSE OP and DT IP were administered to WT control or PF4-iDTR mice. Whole blood, lung and BM were harvested at multiple time points (Figure 4C). Platelets rapidly declined on d1 post-DT and rebounded by d7 (Figure 4C: top right column). CFSE^+^ lung-derived platelets increased on d7 post-DT (Figure 4C: middle left panel) and when normalized to the CFSE^+^ lung Mks, the proportion of lung-derived platelets doubled to 20% (Figure 4C: middle right panel). These data indicated that during acute thrombocytopenia, lung Mks increased their platelet production. DT-mediated depletion targets Mks so we tracked the recovery of lung Mks. On d1 and d2 post-DT, total Mk numbers in both the BM and lung declined relative to controls, and then rebounded by d7 (Figure 4C: bottom left panel). In contrast to some reports of PF4 being expressed in macrophage populations (Pertuy et al., 2015), there was no decline in alveolar or interstitial macrophages on d2-post DT (Supplemental Figure 3A). On d2 post-DT CFSE^+^ lung Mks were reduced and by d7, CFSE^+^ Mks increased relative to controls (Figure 4D: bottom right panel). Because CFSE remained only in the lung tissue, the increase in CFSE^+^ Mks on d7 was from a lung source. These data indicated that thrombocytopenia leads to increased lung-derived platelets.

Antibody-mediated platelet depletion was used as a complementary approach. We observed reduced platelet counts on d1 following anti-Gp1bα antibody, that recovered by d5 (Supplemental Figure 3B, top left panel). However, we did not see an increase in CFSE^+^ platelets (Supplemental Figure 3B, top right panel). When the CFSE^+^ platelets were normalized to CFSE^+^ Mks, we also did not see an increase following antibody treatment (Supplemental Figure 3B, bottom right panel). The reduction in CFSE^+^ platelets following anti-Gp1bα antibody likely occurred due to a reduction in lung Mks, but not BM Mks (Supplemental Figure 3B, bottom left panel).

To test whether lung Mks increase platelet production in a disease context, we used a non-lethal murine malaria model, *Plasmodium yoelli* (PYnL) infection. PYnL results in thrombocytopenia and anemia from about d7-d28 post-infection, at which point the mice clear infection and recover (Vigário et al., 2001). To track lung-derived platelets in infected mice, we gave CFSE OP and PYnL IP on d0 and harvested blood and organs on d7, d14, d21 post-infection, as well as on d40 and d56 when mice had cleared the infection (Figure 4D). We confirmed that the PYnL infection induced a sustained thrombocytopenia (Figure 4D, top). During the chronic thrombocytopenia phase of PYnL, we saw a large increase in circulating CFSE^+^ lung-derived platelets relative to uninfected controls, which normalized during the recovery time points on d40 and d56 (Figure 4D, bottom).

In evaluating the Mk compartments, on d7 there was a decline in BM Mk numbers relative to uninfected controls (Figure 4E: left panel), however, on both d7 and d14 there was an increase in lung Mks relative to uninfected controls (Figure 4E: left panel). On d14 there was a significant decrease in the proportion of CFSE^+^ lung Mks relative to uninfected controls that normalized by 8 weeks after infection (Figure 4E, right panel). These data indicate that there was an influx of lung Mks during infection that came from an extra-pulmonary source. However, after the infection resolved, there was a relative increase in the proportion of lung Mks that were CFSE^+^, suggesting a local lung CFSE^+^ population as the Mk recovery source.

After infection recovery few CFSE^+^ platelets were noted, but CFSE^+^ Mks were still present in the lung. To test whether long-lived/resident lung Mks can be induced to make platelets in PyNL infection, we labeled lung Mks with CFSE dye, waited 3 wks when we reliably find about 10-15% CFSE^+^ Mks in the lung, and infected mice with PYnL (CFSE dye on d0 and infected with PYnL on d21, Figure 4F). 1 wk post-infection (d28 post-CFSE), CFSE^+^ circulating platelets increased in the infected group in comparison with uninfected controls (Figure 4F, left panel), indicating that resident Mks are recruited to make platelets with increased demand. When CFSE^+^ platelets were normalized to CFSE^+^ lung Mks, the infected group’s maximum lung-derived platelet production was about 10% of the circulating platelet pool (Figure 4F, right panel). These data demonstrated that resident Mks produced platelets when challenged with malaria-associated thrombocytopenia.

We also evaluated the ploidy and intravascular/extravascular status of lung Mks on d14 of PYnL infection. There was a significant increase in the proportion of intravascular Mks relative to total lung Mks in PYnL infected mice (Figure 5A top left). This increase was attributed to a significant increase in CFSE^-^ intravascular Mks (Figure 5A top right) and we observed no change in CFSE^+^ intravascular Mks (Figure 5A bottom right). We also saw a decrease in CFSE^+^ extravascular Mks, but similar numbers of CFSE^+^ intravascular Mks at this timepoint (Figure 5A: bottom left), as they had likely left the extravascular space to make platelets. On d14 of PYnL infection, BM Mks had a significant decrease in <4N Mks and a significant increase in 64N+ Mks (Figure 5B). In the spleen, lower ploidy states were depleted, particularly <8N Mks, but 32N and 64N+ Mks were significantly increased (Figure 5B). In contrast, lung Mks remained largely 2-4N, but there was a significant increase in 32N Mks on d14 of PYnL and CFSE^+^ Mks had a significant decline in 2N and 4N Mks at this time point. These data indicated that the extra-pulmonary influx of Mks are low-ploidy, intravascular and platelet producing.

**Figure 5.**
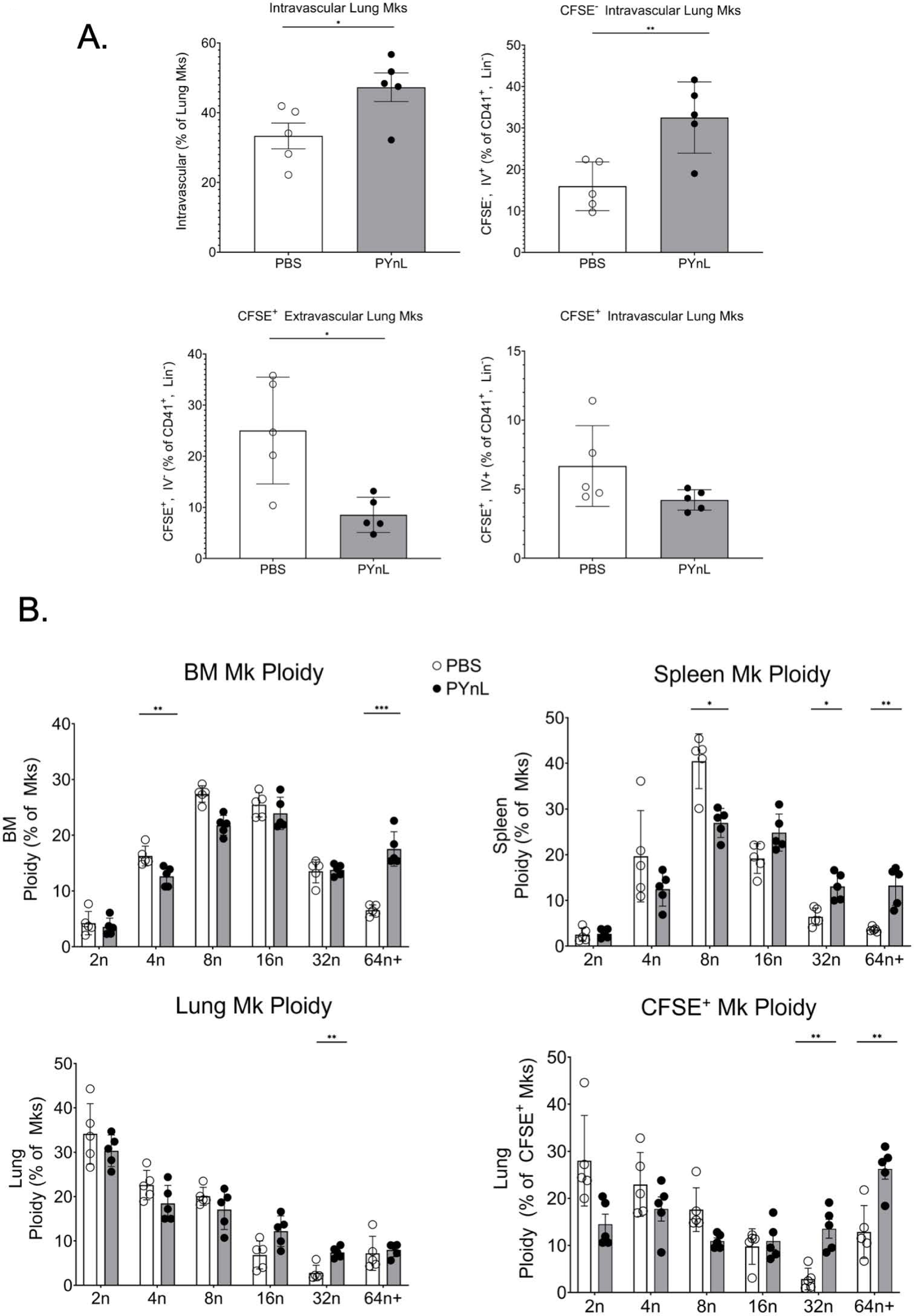
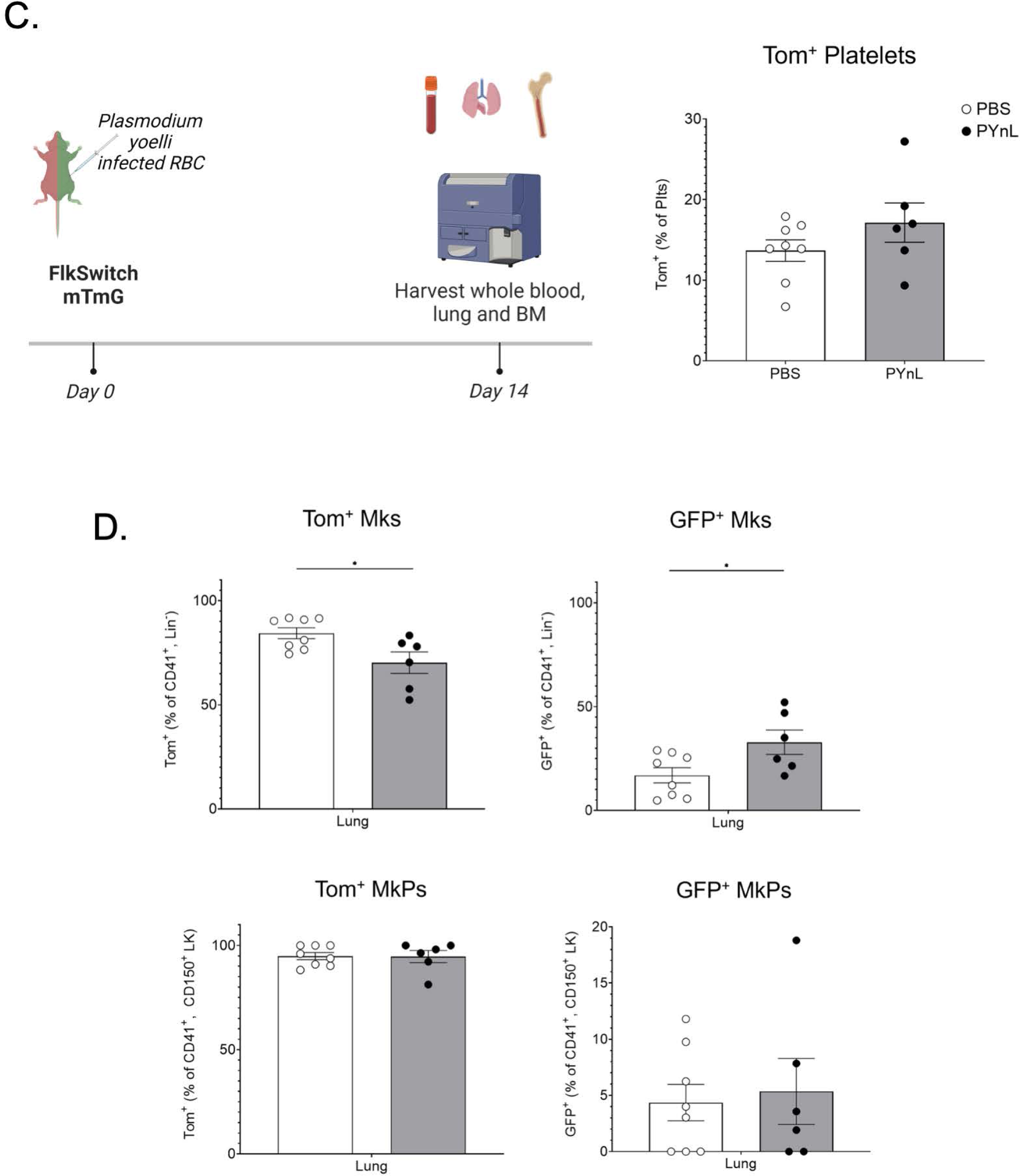
Megakaryocytes migrate to the lung from the BM to respond to increased platelet demand. A) Mice were given CFSE OP, infected with PYnL and on d14 post-infection total and CFSE^+^ and CFSE^-^ intravascular and extravascular Mks were quantified. Post-infection, total intravascular Mks were increased with the increase driven by CFSE^-^ extra-pulmonary Mks. On d14 there was a decrease in CFSE^+^ extra- and intravascular Mks likely driven by their platelet production. B) With PYnL infection there was an increase in higher ploidy CFSE^+^ Mks. C) FlkSwitch mice infected with PYnL had no significant change in Tom^+^/Flt3^-^ platelets on d14 post-infection, but D) the per cent of lung Mks that were Tom^+^/Flt3^-^ slightly declined and GFP^+^/Flt3^+^ increased slightly compared to uninfected controls. There was no change in MkPs indicating an influx of mature Mks from the BM during infection that was still largely Tom^+^/Flt3^-^. (\**p*-value < 0.05, ± SEM; A), C), and D) unpaired two-tailed *t*-tests; B) Multiple *t*-tests with Holm-Sidak multiple comparison correction test)

Because there was an increase in extrapulmonary CFSE^-^ Mks during PYnL infection (Figure 4) we assessed whether Mks that migrate to the lung arise from Flt3^-^ progenitors. FlkSwitch mTmG mice were infected with PYnL and tissue Mks evaluated on d14 post-infection (Figure 5C). There was a slight, but not significant, increase in Tomato^+^ platelets with infection (Figure 5C). The proportion of Tomato^+^/ Flt3^-^ derived Mks in the lung was slightly reduced with a corresponding increase in the proportion of GFP^+^ lung Mks (Figure 5D, top). However, when we evaluated lung Mk progenitors (MkPs), we observed no change in the proportions of GFP^+^ or Tomato^+^ MkP populations (Figure 5D, bottom). These data further imply that increased lung Mks in PYnL infection arise from the BM. Despite the slight increase in GFP^+^ lung Mks at this timepoint, lung Mks were still primarily Flt3^-^/Tomato^+^, indicating that the Flt3^-^/Tomato^+^ Mks have a preference for lung residence, but during hematopoietic stress there may also be a movement of some low ploidy Mks derived from a Flt3^+^ differentiation pathway.

## Discussion

Our data indicates that at least a subpopulation of lung Mks are long-lived, originate via a Flt3-negative pathway from HSCs, and increase platelet production in response to thrombocytopenia. Previous literature assumed that a constant influx of Mks leave the BM, travel through the vasculature, and become ‘stuck’ in the lung making platelets (Lefrançais and Looney, 2019). In contrast, our parabiosis and lung Mk labeling models demonstrated that in a one-month period, only half of lung Mks were replaced by a circulating source. Using CFSE and biotin labeling, we demonstrated that lung Mks can have a lifespan of up to four months while BM Mks turnover within 1-week. These *in vivo* labeling methods validate *in vitro* studies showing that BM Mks live about 5 days (Noetzli et al., 2019). The mechanisms underpinning the prolonged lifespan of lung Mks remains unknown. Our data demonstrated an HSC-dependent, circulatory source for replacement of lung Mks. However, at least some lung Mks remain quiescent or locally proliferate prior to their replacement. The majority of lung Mks are 2N, and perhaps the mechanisms that limit Mk polyploidization also confer their quiescent, yet proliferative, potential. Insights into Mk lifespan may be important for improving current strategies to produce platelets *ex vivo* and will provide important context for studying Mk immune differentiation.

Mounting evidence supports the concept that there is an alternative, direct path from an HSC to Mk that bypasses progenitor stages (Carrelha et al., 2018; Sanjuan-Pla et al., 2013; Shin et al., 2014; Yamamoto et al., 2013). Here, we showed that lung Mks predominantly arise through a direct pathway from a Flt3 negative HSC. In contrast, few BM and spleen Mks (<10%) potentially arise from a direct pathway in baseline conditions. The apparent bias of Flt3^-^ Mks for lung tissue could be due to better fitness of those Mks for the lung environment or perhaps Flt3^-^ Mks have more migratory potential and are more likely to leave the BM, enter the lung, and maintain a longer lifespan. The distinct lineage of lung Mks may also encode their immune modulatory phenotype.

Our methods indicated that lung-resident Mk derived platelet production is about ∼10% of circulating platelets, lower than some previous estimates (Howell and Donahue, 1937; Lefrancais et al., 2017). We cannot however rule out a constant migratory BM-derived Mk source that does not leave the vasculature and significantly contributes to lung-derived platelet production over a shorter lifespan. We also determined that lung resident Mks contribute increased platelets in the settings of both acute and infection-associated thrombocytopenia. The generation of lung Mks via a direct pathway from HSCs would enable lung Mks to serve as a reservoir for platelet production that rapidly responds to thrombocytopenia. Additionally, the immune modulatory phenotype of lung Mks may provide novel mechanisms for detecting thrombocytopenia, particularly during immune challenges.

This work highlights the distinct biology of lung Mks and describes their platelet contribution, particularly during thrombocytopenic states. It remains to be explored if lung Mks generate more immune differentiated platelets. Further research is required to better characterize the role of Mks produced via the direct pathway in all tissue environments. These data add to a growing body of evidence demonstrating the relevance of lung Mks to health and disease.

## Methods

Mice. All mice used were on C57/Bl6 background. Mice not bred in house were purchased from Jackson Laboratory or obtained other collaborators as described below. PF4^iDTR^ mice (Wuescher et al., 2015) were used for megakaryocyte/platelet depletion studies. For depletion, 400 ng of diptheria toxin (Sigma Aldrich D0564-1MG) in sterile water (16 µg/kg) was administered to mice in 100 uL intraperitoneal (IP) injection.

MDS1^CreERT2^ mice were bred with Rosa26-tdTomato as previously described (Zhang et al., 2021). Tamoxifen was given a total of six times spread over two weeks using IP injection for the induction of the fluorescent reporter. Tamoxifen (37.5 mg/kg) was prepared as described (Iturri et al., 2017).

“FlkSwitch” mice were obtained from the University of Utah and have been previously described (Beaudin et al., 2016; Boyer et al., 2011).

CFSE dye oropharyngeal administration: CFSE is a fixable, cell-permeable dye that allows for long-term labeling of cells. CFSE stock (25mM, Thermo Scientific #C1157) was diluted with Iscove’s Media to 8 mM and administered to mice via oropharyngeal (OP) route. Briefly, mice were anesthetized with isoflurane and hung by their teeth on dental floss. The tongue was gently pulled to prevent swallowing. Thirty microliters of 8 mM CFSE was pipetted directly into their oropharyngeal cavity, and mice remained on dental floss for 30-45 seconds to allow for aspiration of CFSE dye. Organs (bone marrow, spleen, and lung) and blood were collected to assess for the presence of CFSE at all time points analyzed. For platelet analyses, CFSE dye was given OP and platelet labeling antibody (Emfret X649) was given via intravenous (IV) route simultaneously. For normalization calculations measuring overall contribution of lung Mks to lung platelets, the following calculation was made.

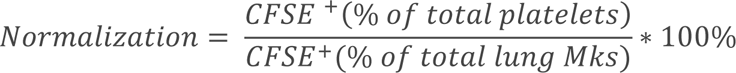

Biotin administration: EZ-link Sulfo-NHS-SS-Biotin (Thermo Scientific #20217) was prepared at 5 mg/mL concentration. Biotin was administered either via oropharyngeal route in 30 microliter volume (as described above) or intravenous as previously described (Nygren and Bryder, 2008). PE-Streptavidin (Biolegend) or BV711-Streptavidin (Biolegend) were used for detection of *in vivo* biotin labeling.

Blood collection, complete blood counts, platelet Isolation and activation: Mice were bled using retro-orbital route into EDTA tubes (Fischer Scientific #NC9990563). Complete blood counts (CBCs) were performed using Abaxis VetScan HM5. Platelets were collected by retro-orbital route into heparanized Tyrodes and isolated as described previously (Cameron et al., 2015). For activation, isolated platelets were incubated with thrombin or U46619 for 10 minutes.

Immune stimuli. For immune stimuli studies, lipopolysaccharide (LPS) or interferon-gamma (IFNγ) were administered to mice via IP route. LPS was given at 1mg/kg for a one-time dose (Sigma-Aldrich #L6259). Recombinant moues IFNγ (R&D Systems, #485MI100/CF) was administered once daily IP at 0.04 mg/kg for three days total.

Plasmodium yoelli (PYnL) was used as a non-lethal chronic infection model of thrombocytopenia. Mice were infected with five hundred thousand infected red blood cells via IP injection and monitored on days (d) 7, 10, 14, 21, 28 and 56 post-infection. For fold change of total Mk counts and CFSE^+^ Mk counts, the following equations were used.

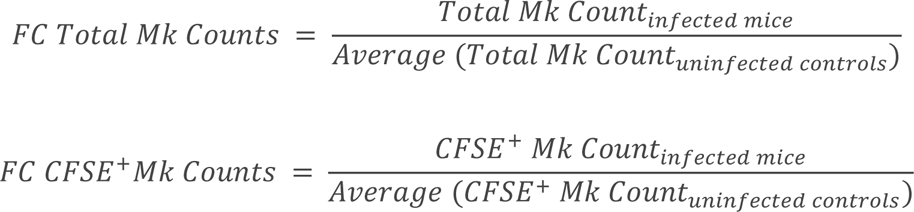

Parabiosis: Parabiosis surgeries were performed surgeries with post-op monitoring as described previously (Kamran et al., 2013; Rodriguez et al., 2022). In brief, 8-week female mice were dosed with CFSE OP 7-10 d prior to surgery. Pre-operative carprofen (5 mg/kg), buprenorphine extended-relase (1 mg/kg), and enrofloxacin (2.5 mg/kg) were administered subcutaneously. Anesthetized WT and mTmG mice were shaved to remove a wide margin of fur at the level of the flank at the intended surgical site of WT and mTmG mice. During surgery mice received matched longitudinal skin incisions from 0.5 cm above the olecranon to 0.5 cm below the knee joint. Subcutaneous tissue was bluntly dissected to expose the olecranon and knee joints of each mouse. Suture was passed around each olecranon and secured with four throws of a square knot. This procedure was repeated at the knee joints. The skin suture was closed using interrupted mattress sutures using 5-0 Vicryl. Interrupted reinforcement sutures were also used. Post-operative monitoring was performed at least twice daily. Multimodal analgesia was continued for at least 3d postoperatively with supplemental soft diet and subcutaneous fluids provided as needed. Analyses of tissues were performed 4 weeks post-surgery.

Single cell suspensions for flow cytometry, ImageStream flow cytometry, and Fluorescence Activated Cell Sorting: For bone marrow single cell suspension, one tibia and one femur were harvested and flushed with isolation buffer using a 20-gauge needle. Isolation buffer consisted of 1 mM EDTA and 2.5% FBS in PBS. Whole lungs were dissected and placed into collagenase type two digestion buffer as described previously (Pariser et al., 2021). All cells were filtered using a 100-micrometer filter and used for downstream analyses. For flow sorting, single cell suspensions were ACK lysed and washed with PBS prior to staining.

Flow cytometry and ImageStream reagents: Antibodies against the following proteins were used: CD41 (MWReg30, Biolegend), Ter119 (Ter119, Biolegend), CD19 (MB19-1, ThermoFisher Scientific), CD3 (17A2, Biolegend), CD11b (M1/70, Biolegend), Gr-1 (RB6-8C5), c-Kit (ACK2, Biolegend), GP1bβ (Emfret, X649), PE-Streptavidin (Biolegend), BV711-Streptavidin (Biolegend), CD62P (RMP-1, Biolegend), Sca1 (D7, Thermofisher Scientific), Flt-3 (A2F10, Thermofisher Scientific), CD48 (HM48-1, Biolegend), CD150 (TC15-12F12.2). Markers used for gating strategies of individual cell types is in Supplemental Table 1.

