## Supplemental Data File for "Lung megakaryocytes are long-lived, arise from Flt3-negative bone marrow cells, and contribute to platelet recovery in thrombocytopenia"

### Supplemental Figures

Supplementary Figure 1

A.

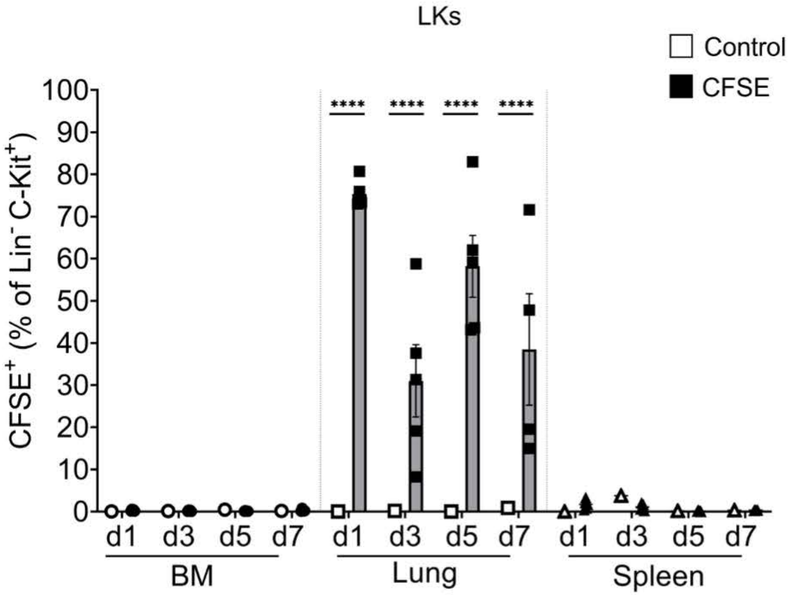


B.

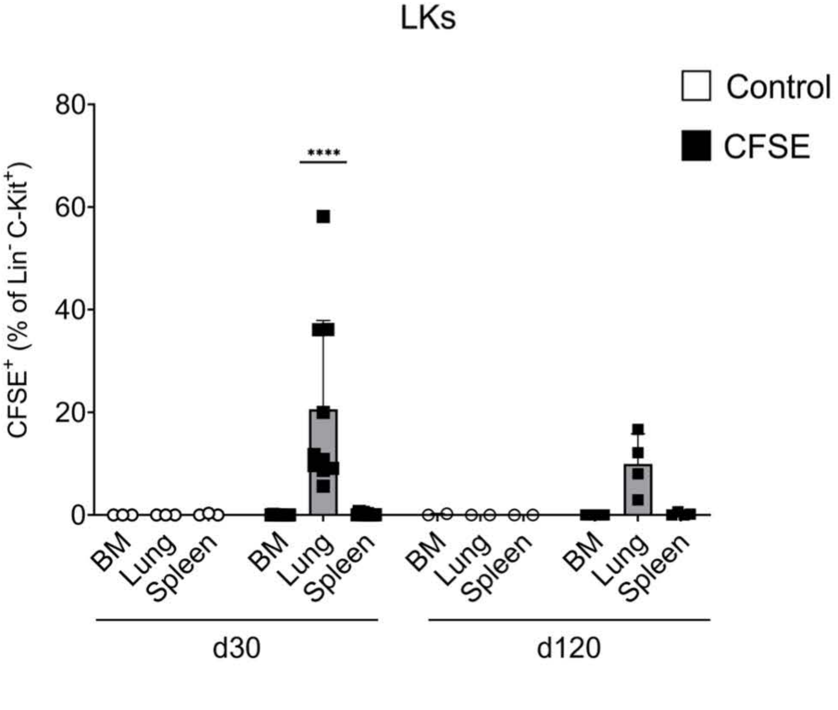


Supplemental Figure 1. Lung HSPCs are Long-Lived. A) CFSE delivered OP to mice specifically labeled lung, but not BM or spleen LKs up to 7d after *in vivo* labeling. B) CFSE labeled lung LKs are present for up to 120d after *in vivo* OP delivery.

Supplementary Figure 2

A.

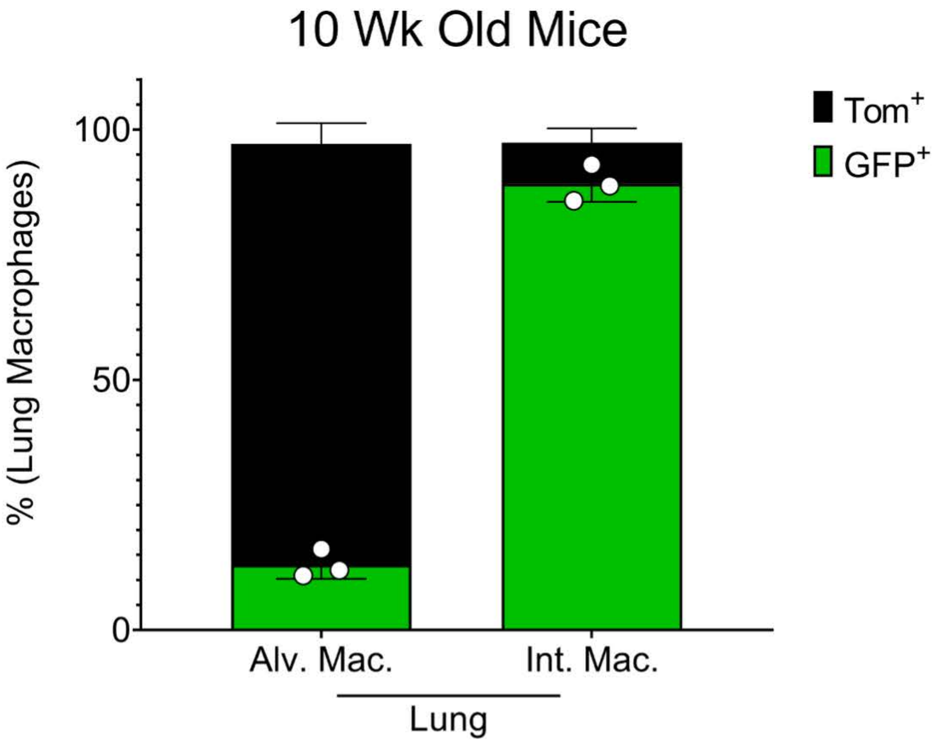


Supplemental Figure 2. As expected, alveolar macrophages are Flt3$^{-}$/Tom+ while interstitial macrophages are Flt3 dependent and GFP$^{+}$.

### Supplementary Figure 3

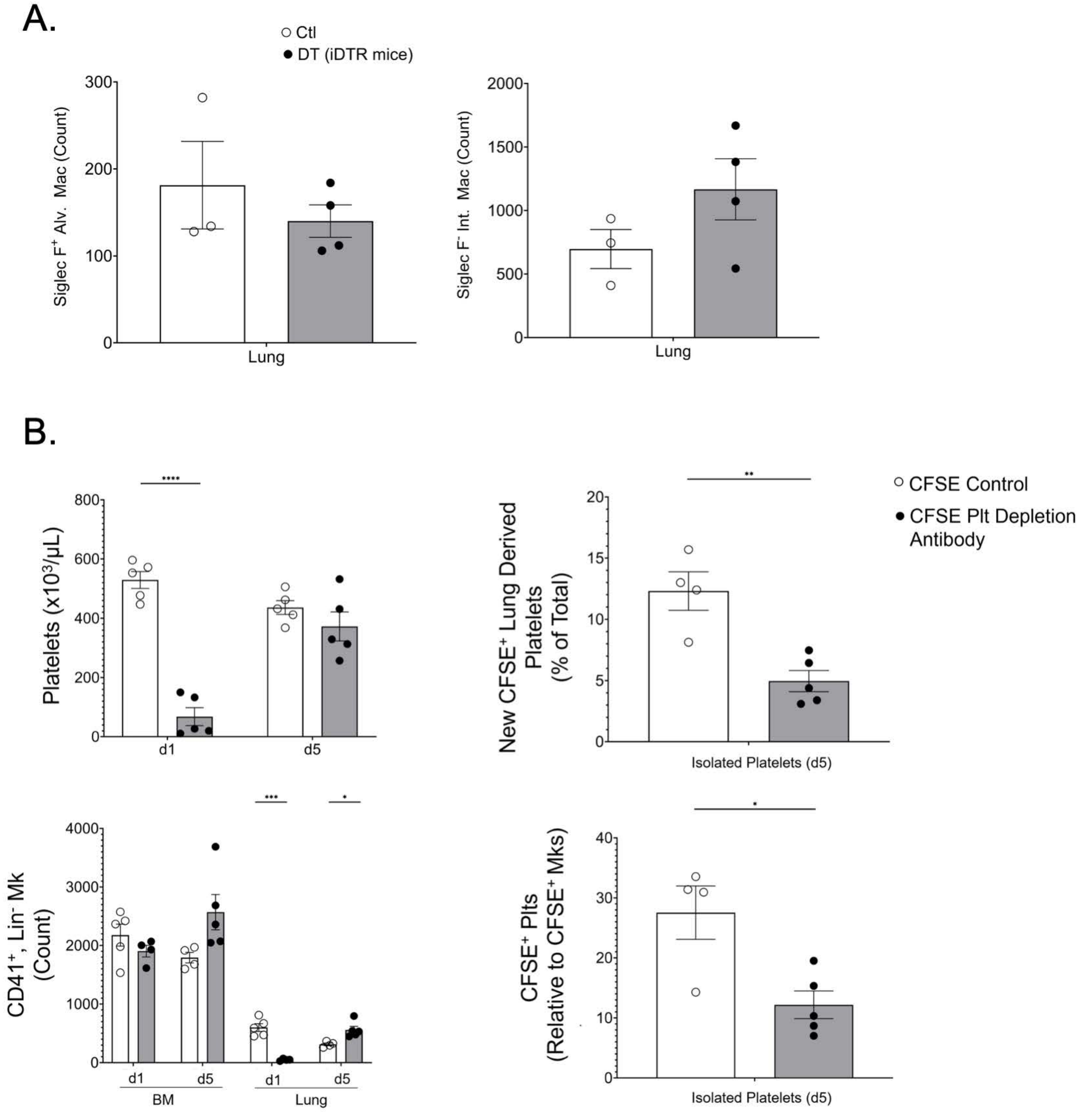


Supplemental Figure 3. A) Neither alveolar nor interstitial macrophage numbers were changed following DT treatment in PF4-iDTR mice. B) Platelet depleting antibody reduced platelet counts and lung but not BM Mks. CFSE+ platelets were also reduced.

Supplemental Table 1. List of markers used for gating strategies for individual cell types.

| Cell type | Markers used for gating strategy |
| --- | --- |
| Platelet | $FSC^{low}$, $SSC^{low}$, $CD41^{+}$ |
| Megakaryocyte (Mk) | $FSC^{intd\text{-}high}$, $SSC^{intd\text{-}high}$, c-$Kit^{-}$, $Lineage^{-}$ (CD3, CD19, CD11b, Gr-1, Ter119), $CD41^{+}$ |
| LSK | $Lineage^{-}$ (CD3, CD19, CD11b, Gr-1, Ter119), $Sca1^{+}$, c-$Kit^{+}$ |
| LK | $Lineage^{-}$ (CD3, CD19, CD11b, Gr-1, Ter119), c-$Kit^{+}$ |
| Long-term hematopoietic stem cell (LT HSC) | $Lineage^{-}$ (CD3, CD19, CD11b, Gr-1, Ter119), $Sca1^{+}$, c-$Kit^{+}$, $Flt3^{-}$, $CD48^{-}$, $CD150^{+}$ |
| Multipotent progenitor 3/4 (MPP3/4) | $Lineage^{-}$ (CD3, CD19, CD11b, Gr-1, Ter119), $Sca1^{+}$, c-$Kit^{+}$, $Flt3^{-}$, $CD48^{+}$, $CD150^{-}$ |
| Multipotent progenitor 2 (MPP2) | $Lineage^{-}$ (CD3, CD19, CD11b, Gr-1, Ter119), $Sca1^{+}$, c-$Kit^{+}$, $Flt3^{-}$, $CD48^{+}$, $CD150^{+}$ |
| Short-term hematopoietic stem cell (ST HSC) | $Lineage^{-}$ (CD3, CD19, CD11b, Gr-1, Ter119), $Sca1^{+}$, c-$Kit^{+}$, $Flt3^{-}$, $CD48^{-}$, $CD150^{-}$ |
| Alveolar Macrophage (AM) | $Ly6G^{-}$, $CD11b^{lo}$, $SiglecF^{hi}$, $CD11c^{hi}$ |
| Interstitial Macrophage (IM) | $Ly6G^{-}$, $CD11b^{+}$, $SiglecF^{-}$, $CD11c^{-}$ |
